# Preliterate symbolic language processing sets the neural stage for learning to read

**DOI:** 10.64898/2026.05.22.726240

**Authors:** Alexia Dalski, Antonia Schulz, Marie Klaes, Max Pirsch, Maria Meinhardt, Agon Ukaj, Laura Faßbender, Vicente Alejandro Aguilera González, Gizem Cetin, Ben de Haas, Yee Lee Shing, Gudrun Schwarzer, Mareike Grotheer

**Author notes:** Corresponding author., Department of Psychology, Philipps-Universität Marburg; Marburg, Germany.

## Abstract

Formal writing is evolutionarily recent, yet the brains of literate adults contain regions - the OTS-words subregions - that respond more strongly to written text than other stimuli. We tested a novel solution to this multi-disciplinary paradox: Does symbolic language processing, which emerged early in human history, lay the neural foundation for reading? In a longitudinal fMRI study, we followed 17 children through their first year of literacy training and related neural responses to text, symbolic language processing, and emerging reading skills over time. We found that middle OTS-words is engaged in symbolic language processing before children learn to read, and that this early engagement predicts later text selectivity and reading ability. These findings suggest that literacy builds on a pre-existing neural scaffold linking vision and language.

## Introduction

Reading ability has a fundamental impact on socioeconomic outlook, societal inclusion and mental health (*1*). Literacy training triggers the emergence of two functional subregions in the left occipito-temporal sulcus (OTS) that respond selectively to texts over other visual stimulus categories (*2*). These subregions - middle OTS-words (mOTS-words, also called VWFA-2) and posterior OTS-words (pOTS-words, also called VWFA-1) - are critical for fluent reading (*3*). However, the mechanisms underlying their emergence remain debated, and three central theories have been proposed: 1) The neuronal recycling hypothesis: This theory proposes that learning to read repurposes neural terrain originally evolved for processing other visual stimuli (*4*). While early evidence suggested that text selectivity recycles face-selective terrain (*5*) more recent work suggests a shift from limb to text selectivity (*6*) during reading acquisition. 2) The theory of graded hemispheric specialization: This theory proposes developmental competition between face and text selectivity in neural terrain that enables high visual acuity, which results in face selectivity becoming increasingly right-lateralized and text selectivity becoming increasingly left-lateralized (*7*). 3) The theory of emergence: This theory proposes that text selectivity develops within previously unspecified cortex and without altering existing visual category selectivity (*8*).

A common theme across these theories is that the OTS-words subregions emerge in neuronal terrain not originally dedicated to reading, as modern writing is a cultural invention that is too recent for dedicated neuronal resources to have emerged through evolutionary mechanisms (*4*). However, even though formal writing systems indeed emerged only ∼5,000 years ago (*9*), it is important to note that humans started producing abstract engravings almost 100.000 years ago (*10*) and hence in a timeframe during which the human brain underwent substantial evolutionary changes (*11*). As such, while formal writing systems recently abstracted away from their iconographic roots (*12*) we may nonetheless have evolved neuronal terrain dedicated to interpreting and combining visual symbols through bio-cultural feedback loops (*13*). Indeed, the ability to recognize visual symbols emerges early in human development, with infants already showing robust understanding of abstract symbols representing common objects (*14*). Here we hypothesize that this early ability to extract meaning from visual symbols could be the scaffold upon which neuronal text selectivity and reading ability is built once literacy training is provided.

To test this hypothesis, we related symbolic language processing, reading ability, and neural text selectivity longitudinally in children (N=17, mean age at study onset ± SD = 6.18 ± 0.39) during the first year of literacy training (i.e. before and after first grade). Two fMRI experiments, embedded within a child-friendly space-narrative (Fig 1A), were collected: In Experiment 1 children viewed texts and other visual stimuli (Fig 1B) to trace the emergence of text selectivity.

**Fig 1:**
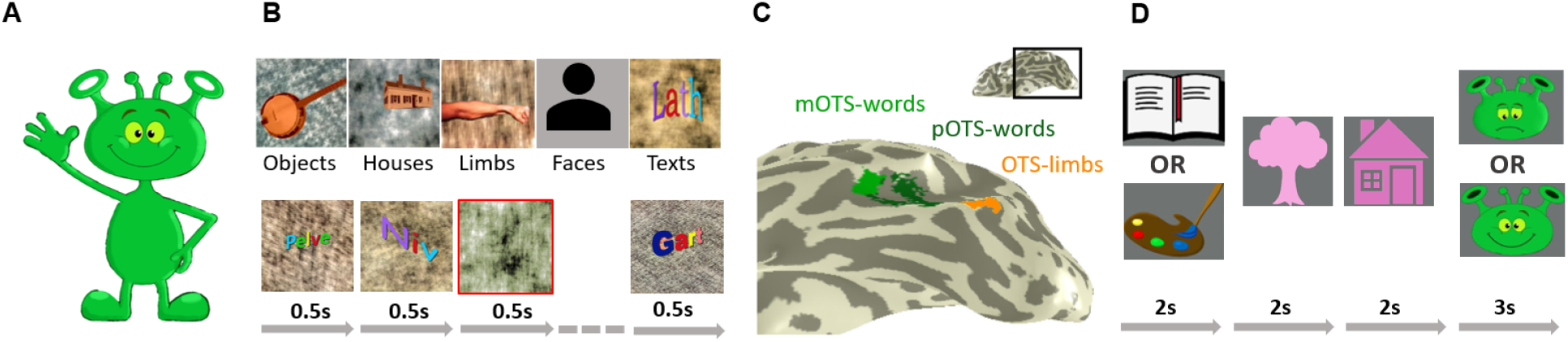
Overview of the experimental design. **A. Child-friendly narrative:** The study was framed as a space adventure in which children assisted Anton the Alien. **B. Experiment 1** : Five visual categories (faces, houses, objects, limbs, texts) were presented in a block design and children performed an oddball task. Top row: example stimuli; Bottom row: example block. Face stimuli in the experiment were photographs of faces presented on the same scrambled background **C. Regions of Interest:** Category selective ROIs were defined from Experiment 1 data collected after first grade and are shown on the inflated cortical surface of an example subject; light green = mOTS-words, dark green = pOTS-words, orange = OTS-limbs. **D. Experiment 2** : Children were presented with pairs of visual symbols and either judged if the two stimuli had the same color hue (color task) or if they formed a meaningful compound noun (symbolic language task). A cue at the beginning of each trial indicated which task children should perform and an Anton-themed answer screen allowed them to provide their answer.

Experiment 1 data collected after first grade was also used to define regions of interest (ROIs) in each child: left mOTS-words and pOTS-words were defined by higher responses to texts than other categories and left OTS-limbs was defined by higher responses to limbs than other categories (T≥3, voxel level, uncorrected, Fig. 1C; all individual ROIs are presented in Supplementary Fig. S1). In Experiment 2 children viewed pairs of visual symbols (Fig 1D, for all stimuli see Supplementary Fig. S2) and were asked to either determine if both symbols had the same color hue (color task) or to judge if the two symbols together form a meaningful German compound noun or not (symbolic language task, e.g. 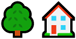(TREE-HOUSE) is meaningful, but 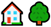(HOUSE-TREE) is not meaningful, for task performance and reaction time see Supplementary Fig. S2). Together, these experiments enabled us to relate children’s emerging reading ability to neural responses evoked by written text and symbolic language processing in the OTS-words subregions.

We find that mOTS-words is engaged in symbolic language processing before first grade and that this early language selectivity predicts changes in stimulus selectivity and reading ability once literacy training is provided. These findings reframe reading acquisition as building on a pre-existing neural scaffold linking language and vision, with broad inter-disciplinary implications.

## Methods

### Participants

Seventeen right-handed children (10 male, 7 female) participated in this longitudinal study with 4 sessions divided over two time-points: Data was acquired once before the start of first grade (2023; mean age = 6.18, years, SD = 0.39) and again after completion of first grade of elementary school (2024; mean age 7.12 years, SD = 0.33). The study was approved by the Ethics Committee of the Medical Faculty of the Philipps University of Marburg (reference 15921). Children were informed verbally in an age-appropriate manner and provided verbal consent. Informed written consent was obtained from the children’s legal guardians. Sessions in which the acquired data did not pass the quality assurance criteria described below were repeated.

### Assessment of reading skill

The children’s reading ability was evaluated at both timepoints using the reading fluency components of the Salzburger Lese- und Rechtschreibtest (SLRT-II), a standardized diagnostic tool for reading and spelling in German-speaking countries (*67*). In these timed components, participants read aloud as many words or pseudowords as possible within one minute to assess their reading fluency. The SLRT-II was administered before and after first grade to track developmental changes in reading fluency. Reading performance is reported as percentile relative to a normative population of children measured after first grade.

### Experimental design

#### Experiment 1

In experiment 1 participants viewed grayscale or colored images of texts (pseudowords), faces, limbs, objects, and houses as in prior work (*68*). Stimuli were presented for 500 ms in blocks of 8 images, each block lasted 4s and 10 blocks per category were presented in each run. Each participant completed four runs (∼5 min each). Participants performed an oddball task, pressing a button whenever a scrambled background without an image appeared. Blocks containing oddball stimuli (∼10% of all blocks) were excluded from analysis. Children successfully detected the oddball stimuli at both timepoints (performance±SE: before first grade: 67±7%; after first grade: 73±5%; reaction times±SE: before first grade: 324±42ms, after first grade: 313±61ms). Performance and reaction times did not differ between categories (performance: F(4, 13) = 0.78, p = 0.57, reaction time: F(4, 13) = 0.88, p = 0.23), did not change across the first year of schooling (performance: F(1,16) = 0.65, p = 0.43; reaction times: F(1,16) = 3.80, p = 0.07), and there was also no time by category interaction (F(4, 13) = 0.91, p = 0.31).

#### Experiment 2

Participants viewed pairs of visual symbols depicting German compound nouns (for all stimuli see Supplementary Fig S1). Symbol pairs either formed meaningful compounds (e.g., “Regen” + “Mantel”, translating to “Rain” + “Coat”) or meaningless combinations (e.g., “Mantel” + “Regen”, translating to “Coat” + “Rain”). While viewing these stimuli, participants performed one of two tasks: In the symbolic language task, participants judged whether the symbol pair represents a meaningful compound noun, and in the color task they judged whether the color hues of the two symbols were identical or slightly different. Pilot experiments run via Amazon Mechanical Turk and locally prior to study onset were used to validate the stimulus set, ensuring that the compound nouns are recognizable for 6 year old children. The pilot experiments were also used to match task difficulty across the language and color tasks.

All images were presented in color on a gray background. Each trial began with a cue (presented for 3 seconds) which indicated which task participants should perform (book pictogram for the symbolic language task, paint palette for the color task). This was followed by the sequential presentation of the two symbols (2 seconds each), that formed the compound noun. Participants indicated their response via a response screen (shown for 3 seconds). Responses were given by selecting a happy (match) or sad (mismatch) Anton the Alien. Only trials in which the children responded were analysed. The inter-stimulus interval between the two symbols was 200 ms. The intervals between the cue and first symbol, the second symbol and the response screen, and the response screen and the subsequent trial were each jittered randomly to either 1 or 2 seconds. Participants completed six runs (∼5 min each), with 16 trials per run. Task and stimulus conditions were balanced such that each compound noun appeared equally often across conditions. Conditions were pseudorandomized in such a way that the task switched only every four trials, with the starting task counterbalanced across runs. This pseudorandomization was implemented to increase child-friendlyness, as task switching is challenging for young children.

In experiment 2, performance (% correct ±SE; Supplementary Fig. S2A) exceeded chance-level at both timepoints in both the symbolic language task (before first grade: 64±5%, t(16) = 2.58, p = 0.02; after first grade: 78±4%, t(16) = 6.98, p < 0.0001) and the color task (before first grade: 60±5%, t(16) = 2.27, p = 0.04; after first grade: 78±5%, t(16) = 5.79, p < 0.0001). Performance improved significantly across the first year of schooling (main effect of time; F(1,16) = 7.96, p = 0.012), did not differ between the tasks (no main effect of task (F(1,16) = 0.40, p = 0.54), and showed no task × time interaction (F(1,16) = 0.73, p = 0.41). Reaction times (Supplementary Fig. S2B) did not change across the first year of schooling (mean±SE before first grade: language task: 1088±77ms, color task: 1012±77ms; mean±SE after first grade: language task: 1098±70ms, color task: 1124±60ms, no main effect of time F(1,16) = 0.58, p = 0.46), did not differ between tasks (no main effect of task: F(1,16) = 0.39, p = 0.54) and showed no task × time interaction (F(1,16) = 2.60, p = 0.13).

### Custom procedures for neuroimaging in children

Collecting neuroimaging data from children is most successful with customized procedures and we have implemented several important steps to ensure participant well-being and data quality. First, prior to scanning, children and their families were familiarized with the scanner environment through a child-friendly instructional video. Second, a dedicated additional staff member accompanied the child and was present in the scanner room at all times to ensure the child’s well-being and to motivate the child. Third, all scan sessions were kept to a maximum of 30 minutes to reduce fatigue and boredom. Fourth, both experimental paradigms were designed to be age-appropriate: Experiment 1: This experiment paradigm has already been used successfully in developmental populations (*6*) and is particularly child-friendly because: i) The oddball detection task is comparatively simple and ii) The experiment takes only 20 minutes to complete. Experiment 2: We designed this paradigm to be age appropriate by: i) Using German compound nouns that are familiar to 6-year old children, ii) Presenting the stimuli and the answer screen for longer durations, and iii) Using a blocked task structure, in which the task switched only every four trials, since task-switching is challenging for children at this age. To ensure task comprehension, children completed a trial run of each experiment before entering the MRI. Both experiments were also embedded in a playful narrative in which “Anton the Alien” is visiting in his spaceship - the MRI scanner, decorated with space-themed stickers.

### Neuroimaging data acquisition

All measurements were conducted at the Bender Institute for Neuroimaging (BION), Justus Liebig University Giessen, on a Siemens Magnetom Prisma 3T scanner with a 64-channel head-neck coil. Functional images were acquired using a T2*-weighted gradient-echo EPI sequence (TR = 1000 ms, TE = 30 ms, flip angle = 59°). Each volume comprised 42 axial slices covering the occipitotemporal cortex and parts of the frontal cortex (voxel size = 2.5 × 2.5 × 2.5 mm^3^; FoV = 192 mm). High-resolution anatomical images were collected with a T1-weighted BRAVO sequence (TR = 450 ms, TE = 3.53 ms, flip angle = 12°, voxel size = 1 × 1 × 1 mm^3^; FoV = 240 mm).

### Neuroimaging data preprocessing

Anatomical volumes were segmented into gray and white matter using FreeSurfer (http://surfer.nmr.mgh.harvard.edu/). As children’s brains may still undergo developmental changes, we used FreeSurfer’s longitudinal pipeline to ensure consistent surface reconstruction across timepoints. For each child, scans from before and after first grade were reconstructed separately, then combined into an unbiased within-subject template. Each timepoint was subsequently reprocessed relative to this template, ensuring alignment in a common reference frame. Segmentations were manually corrected in ITKGray.

Functional data were analyzed with the mrVista toolbox (http://github.com/vistalab) for Matlab. Functional runs were motion-corrected within and across runs, aligned to the participant’s anatomy, and analyzed in native space without spatial smoothing. Voxel time courses were high-pass filtered at 0.05 Hz. For each experiment, a design matrix was convolved with the SPM hemodynamic response function (http://www.fil.ion.ucl.ac.uk/spm) to generate predictors, and regularized response coefficients (betas) were estimated with a general linear model (GLM). No spatial smoothing was applied, and group-based analyses were avoided to reduce spurious overlap (*17*).

### Neuroimaging data quality assurance

Several steps were taken to reduce motion at the time of data collection including: i) detailed and child-friendly instructions and narrative, ii) constant monitoring of the child by a dedicated staff member inside the scanner room, iii) short scanning sessions limited to <30 minutes. During data processing, motion thresholds matched those we used for adults in prior work (*18*): Functional data in which motion exceeding 3 voxels within a run were excluded (17 out of 272 runs) and sessions were re-collected when fewer than 3 usable runs remained (10 out of 68 sessions). Functional data in which between-run motion exceeded 3.5 voxels were similarly re-collected (1 out of 68 sessions).

### Regions of interest definition (ROIs)

Functional regions of interest (ROIs) were defined from experiment 1 data collected after first grade. Three ROIs (T=3; uncorrected) were defined in the left occipito-temporal sulcus (OTS) in each participant’s native space based on both functional and anatomical criteria (all ROIs are presented in Supplementary Fig. S3):

- mOTS-words: A region in the middle OTS that shows stronger responses to texts than other visual stimulus categories (limbs, faces, houses, objects).
- pOTS-words: A region in the posterior OTS that shows stronger responses to texts than other categories (limbs, faces, houses, objects).
- OTS-limbs: A region in the OTS that is adjacent to mOTS-words and pOTS-words but shows stronger responses to limbs than other categories (texts, faces, houses, objects) and serves as a control region.

We defined mOTS-words (mean size: 218 mm^3^, SD = 206 mm^3^) and pOTS-words (mean size: 465 mm^3^, SD = 402 mm^3^) in all 17 participants. For 5 participants experiment 1 was collected twice after first grade, to ensure that OTS-words could be defined. OTS-limbs could be defined in 16 out of 17 participants (mean size: 390 mm^3^, SD = 280 mm^3^). For the split-half approach (odd runs only), we were able to define left mOTS-words in 15 out of 17 participants (mean size: 178 mm^3^, SD = 201 mm^3^), left pOTS-words in 15 out of 17 participants (mean size: 421 mm^3^, SD = 358 mm^3^), and left OTS-limbs in 15 out of 17 participants (mean size: 345 mm^3^, SD = 303 mm^3^).

### Calculation of Selectivity Indices

Estimated betas from the three ROIs (mOTS-words, pOTS-words, and OTS-limbs) reflecting response magnitude in each condition in each experiment are presented in Supplementary Figs. S4 and S5. Beta weights indicate the overall magnitude of a brain region’s response to a given condition, but do not reflect a region’s response to one condition relative to others. To quantify this relative preference, we computed selectivity indices (SIs) at each timepoint and for each experiment from the betas as in prior work (*19*)). SIs range from -1 to +1, where positive values indicate stronger responses to the current condition relative to all others, values near zero indicate no preference, and negative values indicate stronger responses to the other condition(s). The SIs allowed us to track how our ROIs functional tuning changes across the first year of schooling and enabled us to relate changes in functional tuning to changes in reading skill.

For experiment 1, SIs were computed as: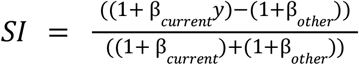

Where β_*current*_ is the beta estimate for the current category of interest and β_*other*_ is the average beta of all remaining categories.

For experiment 2, SIs were computed as: 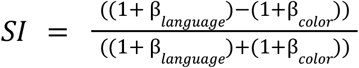

Where β_*language*_ refers to the symbolic language and β_*other*_ refers to the color task. Because there are only two task conditions (symbolic language and color), a single SI per participant and timepoint captures the full contrast between them: a positive SI indicates a preference for the symbolic language task, a negative SI indicates a preference for the color task.

### Statistical analyses

#### Behavioral measures

Reading fluency in the Salzburger Lese- und Rechtschreibtest II (SLRT-II, real- and pseudo-word reading, measured as number of words read correctly per minute and converted to percentile relative to a normative population) was compared before and after first grade using a paired t-test (Fig. 4A).

Behavioral responses (percent correct responses and reaction times) in the fMRI experiments were evaluated with rmANOVAs that used category and time (experiment 1) or task and time (experiment 2) as factors. In experiment 2 we further used one-sample t-tests to assess whether accuracy exceeded 50% chance level.

#### fMRI responses

Betas weights from each ROI were assessed using rmANOVAs with category and time as factors (experiment 1, Supplementary Figure S4) or with task and time as factors (Experiment 2, Supplementary Figure S5). Where rmANOVAs revealed significant main effects or interactions, post-hoc pairwise comparisons were performed.

Next, betas were converted to selectivity indices (SIs, as described above) and SIs were assessed. In experiment 1, rmANOVAs were fitted with the factors category and time (Fig. 2). In experiment 2, rmANOVAs were fitted with the factor time and one-sample t-tests were used to test SIs against zero separately at each timepoint (Fig. 3A-C). Where rmANOVAs revealed significant main effects or interactions, post-hoc pairwise comparisons were performed.

**Fig. 2.**
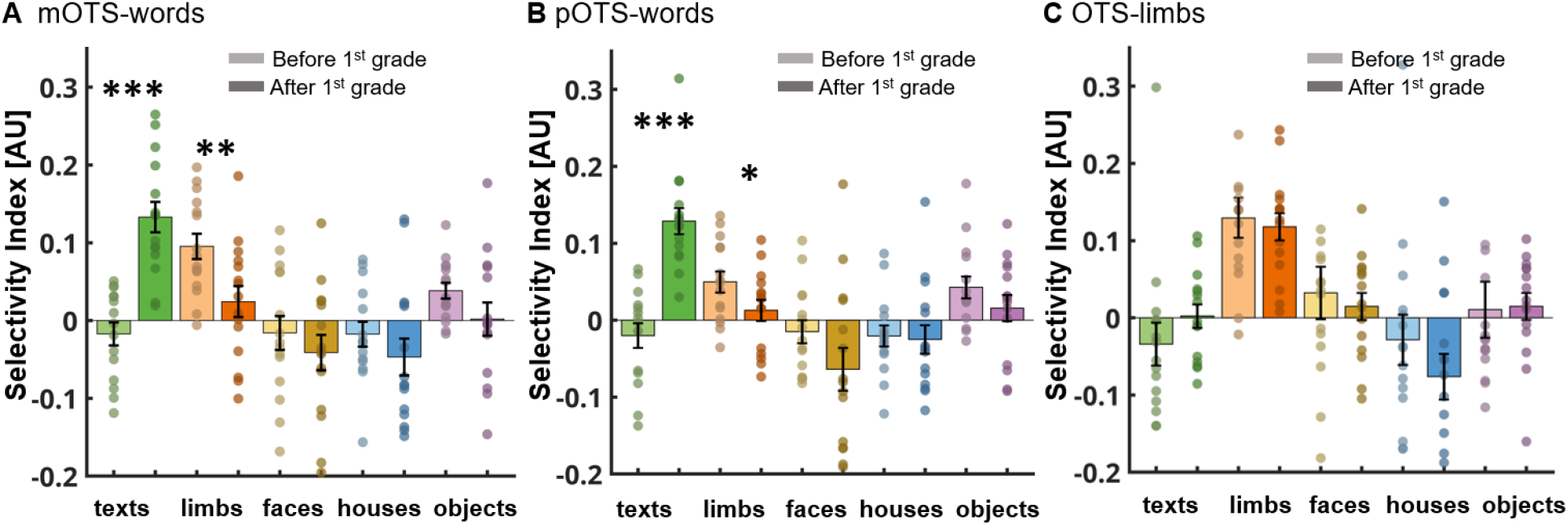
During the first year of literacy training, selectivity for texts increases and selectivity for limbs decreases in the OTS-words subregions. (A-C) Mean selectivity indices (± SEM) for five visual categories (texts, limbs, faces, houses, and objects) are shown before (lighter shades) and after (darker shades) first grade in three ROIs: mOTS-words (A), pOTS-words (B) and OTS-limbs (C). Positive SI values indicate stronger responses to the current category relative to other categories. Asterisks indicate significant changes in category selectivity between timepoints (\*\*\**p*<0.001, **p<0.01, *p<0.05), circles indicate individual subject data.

**Fig 3.**
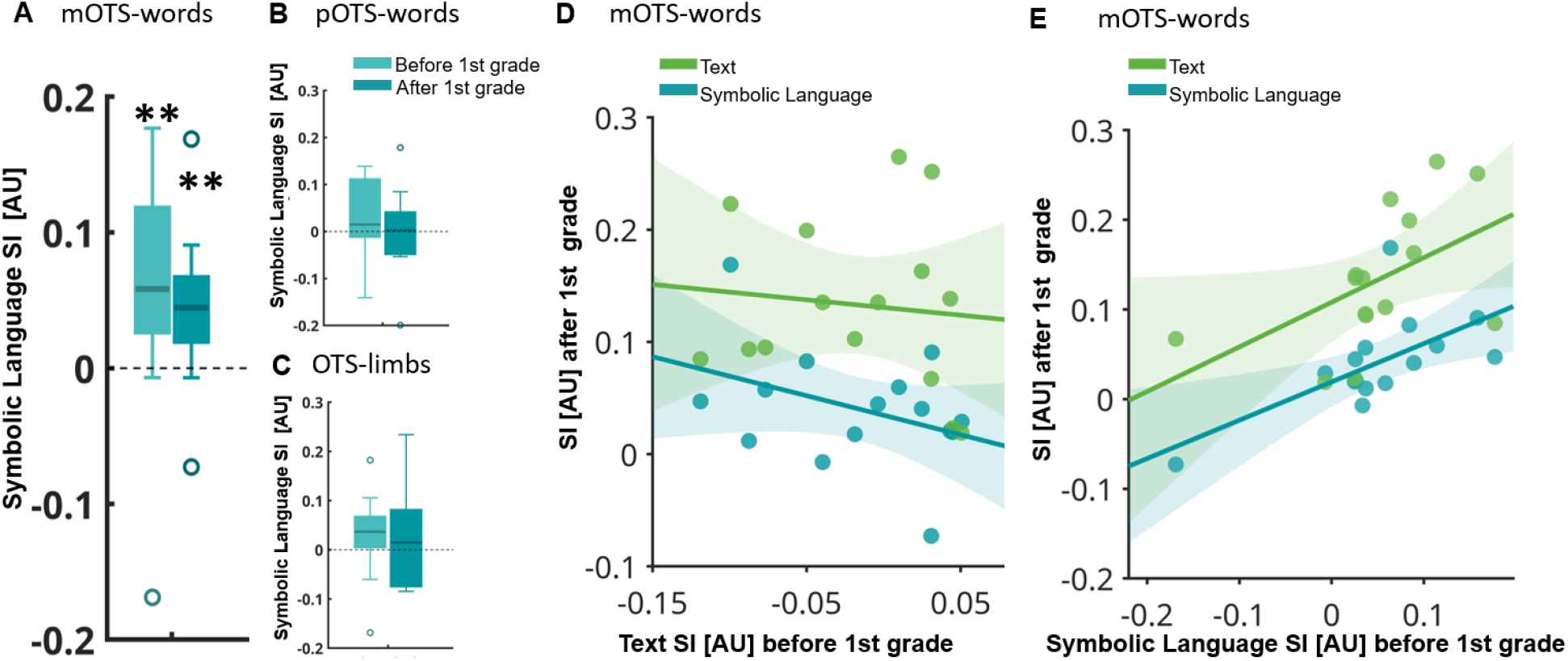
Symbolic language processing in mOTS-words precedes schooling and predicts emerging text selectivity. (A-C) Selectivity indices for language processing are shown before (lighter shades) and after (darker shades) first grade in three ROIs: mOTS-words (A), pOTS-words (B) and OTS-limbs (C). Boxes show the interquartile range (IQR), with the horizontal line indicating the median; whiskers extend to the most extreme values within 1.5 × IQR of the first and third quartiles; circles indicate outliers. Positive SI values indicate stronger responses in the language task than the color task. Asterisks indicate t-tests against 0 for each timepoint; \*\**p*<0.01. (D-E) Prediction of mOTS-words neural selectivity after first grade from mOTS-words text selectivity (D) or symbolic language selectivity (E) before first grade. Text selectivity before first grade does not predict future text selectivity or symbolic language processing, but symbolic language selectivity before first grade predicts both future text selectivity and future language selectivity. Each circle is an individual, lines show least-squares regression fits, shaded regions show 95 % confidence intervals.

#### Correlations of selectivity indices across experiments and timepoints

We related SIs across experiments and the two time points using Pearson correlations (reported as r^2^ and p values) computed separately for each ROI. First, we tested whether text selectivity before first grade relates to text selectivity or symbolic language processing after first grade, or to the increase in text selectivity during schooling (Fig. 3D). Second, we tested whether symbolic language processing before first grade relates to symbolic language processing after first grade, text selectivity after first grade, or each individual’s increase in text selectivity across the first year of schooling (Fig. 3E).

#### Correlations of selectivity indices with behavioral reading performance

To assess the relationship between neural selectivities and reading fluency after the first year of reading instruction, we computed Pearson correlations (reported as r^2^ and p values) relating text selectivity and symbolic language selectivity at both timepoints to real-word reading fluency (Fig. 4B,C) as well as pseudo-word reading fluency (Supplementary Fig. S6) after first grade (SLRT-II percentile ranks), separately for each ROI.

**Fig. 4.**
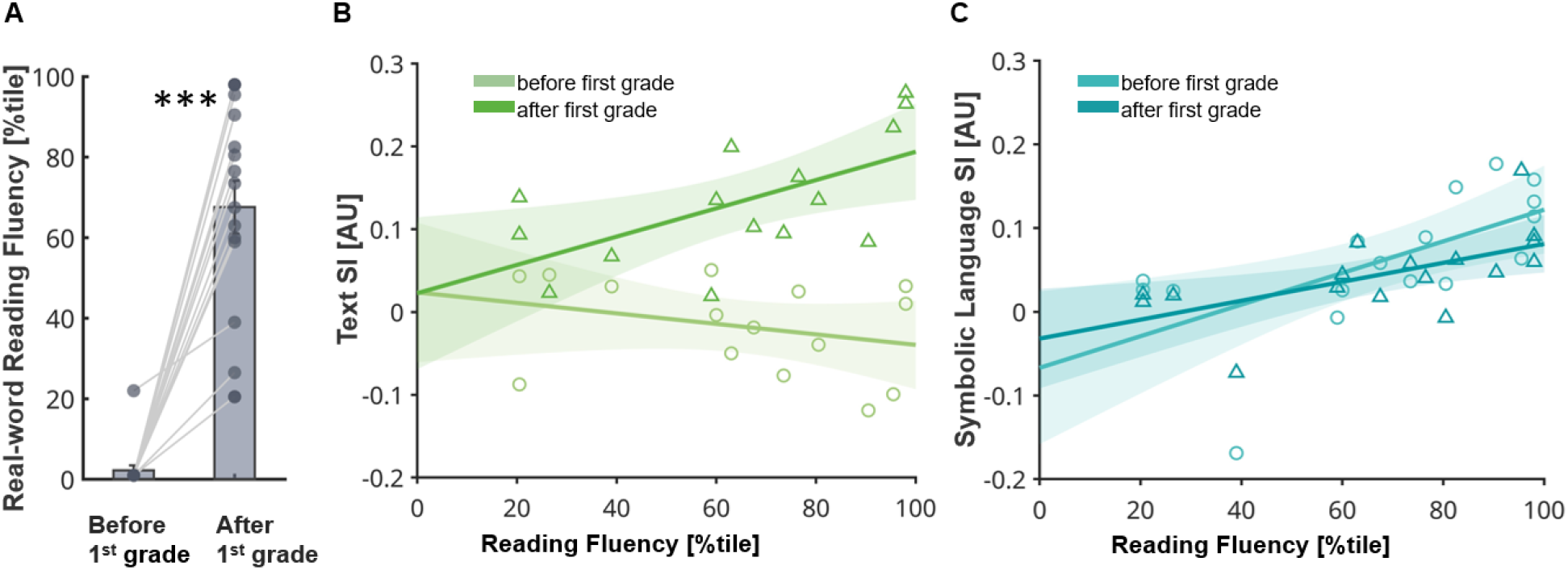
Preliterate symbolic language processing in mOTS-words predicts future reading fluency. (A) Children’s reading ability (SLRT-II, real-word reading fluency) increased during the first year of schooling (paired t-test, p<0.0001). Bars show mean percentile (± SEM), each circle is an individual, lines connect the two measurements of the same child. (B) Individual children’s (N=15) reading fluency after first grade correlates with mOTS-words text selectivity after first grade (triangles, dark green) but not before first grade (circles, light green). (C) Individual children’s (N=17) reading fluency after first grade correlates with mOTS-words symbolic language selectivity after first grade (triangles, dark cyan) and strikingly it is predicted by mOTS-words language selectivity before first grade (circles, light cyan). In (B,C) each triangle or circle is an individual, lines show least-squares regression fits, shaded regions show 95 % confidence intervals.

## Results

### Neural text-selectivity emerges during the first year of literacy training

First, we evaluated changes in visual category-selectivity using experiment 1 data. General linear models were used to estimate beta weights for responses to all stimulus categories and betas were extracted from the three ROIs as defined after schooling. To avoid circularity when evaluating the second timepoint, the data was split into 2 independent samples - one sample was used to define ROIs and the other sample was used to evaluate their neural response profile. Only children (N=15) in whom all ROIs could be defined in this way were included. To quantify visual stimulus selectivity of our ROIs and track how it changes with schooling, we computed selectivity indices (SIs):

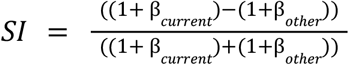

Where β_*current*_ is the beta estimate for the current category of interest and β_*other*_ is the average beta of all remaining categories.

Both mOTS-words and pOTS-words SIs showed significant stimulus selectivity that changed over the first year of schooling (rmANOVA; main effect of category: mOTS-words: F(4,11)=9.81, p<0.0001; pOTS-words: F(4,11)=9.27, p<0.0001; category×time interaction: mOTS-words: F(1,14)=11.71, p<0.0001; pOTS-words: F(4,11)=13.51, p<0.0001). Post-hoc analyses of the OTS-words subregions revealed highest SI for limbs before first grade in mOTS-words but not pOTS-words (mOTS-words: responses to limbs higher than all other categories: p<0.05 for all pairwise comparisons, pOTS-words no preference for any category: p>0.05 for all pairwise comparisons). Both subregions further showed highest SI for texts after first grade (mOTS-words: p<0.05 for all pairwise comparisons, pOTS-words: p<0.05 for all pairwise comparisons), as well as a significant increase in text selectivity (mOTS-words: p<0.0001; pOTS-words: p<0.0001), with a concurrent decrease in limb selectivity (mOTS-words: p=0.002; pOTS-words: p=0.04) during the first year of schooling. In contrast, while OTS-limbs SIs also showed significant stimulus selectivity (rmANOVA; main effect of category: F(4,11)=16.55, p<0.0001), this stimulus selectivity did not change during the first year of schooling (category×time interaction not significant: F(4,11)=1.65, p=0.17). Post-hoc analyses of OTS-limbs revealed highest SIs for limbs (p<0.05 for all pairwise comparisons).

Repeated-measures ANOVAs of raw beta values corroborate these findings and similarly show changing category selectivity in the OTS-words subregions (mOTS-words: main effect of category: F(4,11)=7.63, p<0.0001, category×time interaction: F(4,11)=8.20, p<0.0001; pOTS-words: main effect of category: F(4,11)=10.98, p<0.0001, categoryxtime interaction: F(4,11)=16.05, p<0.0001) but stable category selectivity in OTS-limbs (main effect of category: F(4,11)=8.59, p<0.0001, no category×time interaction: F(4,11) = 0.93, p = 0.45) (Supplementary Fig. S4).

### Symbolic language processing precedes literacy training in mOTS-words

Next, we explored each ROIs engagement in symbolic language processing by computing the Sis for a language task over a color task 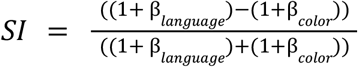 performed on visual symbols in experiment 2. To compute the SIs, betas were extracted from the three ROIs as defined from experiment 1 data after schooling (N=17). Strikingly, mOTS-words showed a preference for symbolic language processing that was stable over time (rmANOVA; no effect of time, F(1,16)=1.23, p=0.28) and observable both before and after the first year of schooling (t-test against 0; before first grade: t(16)=3.12, p=0.007; after first grade: t(16)=3.63, p=0.002) (Fig. 3A). In contrast, pOTS-words and OTS-limbs showed no language selectivity either before or after the first year of schooling (t-test against 0; pOTS-words: before first grade: t(16)=1.85, p=0.08, after first grade: t(16)=0.40, p=0.70; OTS-limbs: before first grade: t(15)=1.55, p=0.14, after first grade: t(15)=0.17, p=0.87) (Fig. 3B,C). In pOTS-words and OTS-limbs, symbolic language selectivity also did not change over time (rmANOVA; no main effect of time: pOTS-words: F(1,16)=0.82, p = 0.38; OTS-limbs: F(1,15)=0.14, p=0.71).

Repeated-measures ANOVAs of raw beta values corroborate these findings and similarly show a preference for symbolic language processing in mOTS-words (main effect of task: F(1,16)=11.34, p=0.004) that does not change over time (no taskxtime interaction: F(1,16)=1.49, p=0.24) (Supplementary Fig. S5) and is not detectable in either pOTS-words or OTS-limbs (no main effect of task: pOTS-words: F(1,16)=2.15, p=0.16, OTS-limbs: F(1,16)=0.03, p=0.88).

### Symbolic language processing predicts future text-selectivity in the OTS-words subregions

In the next step, in all ROIs, we related neural selectivities across time and between the two experiments. Text selectivity before first grade did not predict text selectivity after first grade in any of our ROIs (Pearson correlations; mOTS-words: r^2^=0.01, p=0.71; pOTS-words: r^2^=0.01, p=0.70; OTS-limbs: r^2^=0.13, p=0.18) (Fig. 3D-green). Similarly text selectivity before first grade also did not predict symbolic language selectivity after first grade in any of our ROIs (mOTS-words: r^2^=0.15, p=0.16; pOTS-words: r^2^=0.01, p = 0.77; OTS-limbs: r^2^=0.08, p=0.31) (Fig. 3D-cyan). Symbolic language processing before first grade, on the other hand, did predict symbolic language selectivity after first grade in mOTS-words (r^2^=0.43, p=0.004) (Fig 3E-cyan), but not pOTS-words (r^2^=0.00, p=0.87) or OTS-limbs (r^2^=0.03, p=0.51). Most strikingly, symbolic language processing before first grade predicted text selectivity after first grade in both mOTS-words (r^2^=0.27, p=0.05) (Fig. 3E-green) and pOTS-words (r^2^=0.37, p=0.02), but not in OTS-limbs (r^2^=0.12, p=0.20). Finally, symbolic language processing before first grade also predicted each individual’s increase in text selectivity during the first year of literacy training in mOTS-words (r^2^=0.35, p=0.02) and pOTS-words (r^2^=0.30, p=0.04) but not OTS-limbs (r^2^=0.15, p=0.16).

### Preliterate symbolic language processing in mOTS-words predicts future reading skill

As expected, children’s reading ability (Salzburger Lese- und Rechtschreibtest II (SLRT-II), real-word reading fluency as assessed by number of words read correctly per minute, reported as percentiles relative to normal population measured after first grade) increased during the first year of schooling (Fig. 4A, paired t-test: t(16) = 9.35, p<0.0001). Individual children’s real-word reading fluency after first grade correlated with their mOTS-words’ text selectivity after first grade (r^2^=0.37, p=0.02), but not with their mOTS-words text selectivity before first grade (r^2^=0.09 p=0.28) (Fig. 4B). No correlation between reading fluency after first grade and text selectivity was observed in pOTS-words or OTS-limbs (all r^2^<0.14, all ps>0.16). Individual children’s reading fluency after first grade was also correlated with their mOTS-words symbolic language selectivity after first grade (r^2^=0.37, p=0.01) and strikingly it was predicted by their mOTS-words language selectivity before first grade (r^2^=0.41, p=0.006) (Fig. 4C).

A similar pattern was observed for pseudo-word reading fluency: mOTS-words symbolic language selectivity, but not text selectivity, before first grade predicted pseudo-word reading fluency after first grade (language selectivity: r^2^=0.43, p=0.004; text selectivity: r^2^=0.12, p=0.20; Fig. S6). No correlation between pseudo-word reading fluency after first grade and symbolic language processing or text selectivity was observed in pOTS-words or OTS-limbs (symbolic language: all r^2^<0.10, all ps>0.20, text: all r^2^<0.09, all ps>0.28).

## Discussion

In this longitudinal study, we evaluated the neural mechanisms that enable children to become fluent readers. Focusing on the OTS-words subregions - key brain regions that are critical for reading (*3, 20*) - we find that mOTS-words is engaged in symbolic language processing before first grade and that this early engagement predicts changes in neural text selectivity and reading ability during literacy training.

Our findings suggest that mOTS-words is involved in symbolic language processing prior to and independently of schooling. The region may hence represent a central hub bridging vision and language that is recruited during literacy training. Literacy training, in turn, shapes the region’s visual selectivity for text to facilitate reading fluency. This view aligns with prior studies on the phonological basis of reading which similarly show that language-related functional and structural neural architecture early in life shape later phonological ability and reading skills across individuals (*21*–*23*). Further evidence in support of this theory comes from connectivity studies showing that OTS-words is both structurally (*24*–*27*) and functionally (*28, 29*) connected not only to visual, but also to language regions. Remarkably, this connectivity pattern is so consistent, that it can be used to predict where the region is located in literate adults (*30*) and even where it will emerge in preliterate children once they learn how to read (*31, 32*). When incorporating this privileged connectivity pattern, the view of OTS-words as a vision-to-language bridge, also explains why mOTS-words emerges in a similar cortical location across individuals and in a large variety of writing systems (*33*). Moreover, this view aligns with amounting evidence showing increased OTS-words responses when literate adults engage in language-related tasks, even when these tasks are performed on stimuli other than text (*18, 25, 34, 35*).

From prior theories on the emergence of OTS-words, our data best align with the revised neuronal recycling hypothesis, which posits that during reading acquisition neural resources originally dedicated to processing limbs become recycled for processing text (*6, 36*). Indeed, similar to prior work (*6*), we find that increases in text selectivity during first grade are accompanied by decreases in limb selectivity in the OTS-words subregions. However, while this shift from limb to text selectivity has previously been explained in the context of changes in children’s visual diet (*6, 36, 37*), i.e. the amount of time children spend looking at limbs and at texts (*38*), our data which incorporates symbolic language processing offers an alternative mechanistic explanation: as younger children rely on their hands for communication to a greater extent than older children with more developed language skills (*39*), the observed decrease in limb selectivity with a concurrent increase in text selectivity may be related to changes in the degree to which vision-language mapping is needed for these stimulus classes. This alternative mechanistic explanation can also account for the observation that OTS-words is recruited for Braille-reading in congenitally blind individuals (*40, 41*) in whom no changes in visual diet occurs.

Interestingly, while we find a shift from limb to text selectivity during schooling in both mOTS-words and pOTS-words, only mOTS-words is engaged in symbolic language processing prior to first grade. These differences suggest that the OTS-words subregions may emerge through distinct developmental mechanisms, which aligns with prior work differentiating the two sub-regions: First, mOTS-words and pOTS-words are located in distinct cytoarchitectonic areas (*42, 43*). Second, functional (*44*) and structural (*25, 26, 30*) connectivity studies showed that while mOTS-words is preferentially connected to language regions, pOTS-words is preferentially connected to other visually-driven regions. Third, the stimulus preferences of the subregions differ with pOTS-words being sensitive to lower-level visual properties, and mOTS-words being sensitive to higher-level lexical properties of the presented stimuli (*26, 45, 46*). These dissociations reveal subregion-specific anatomy and computational roles and thereby emphasize the need to evaluate the two subregions separately.

One critical aspect that distinguishes our findings from prior theories is the observation that mOTS-words is involved in symbolic language processing before literacy training, making it predisposed for supporting reading before children learn to do so. Typically it is argued that reading is a too novel cultural invention to allow for such a predisposition, as dedicated neural architecture for reading could not have been laid out through evolutionary mechanisms (*4*). However, even though formal writing systems indeed emerged only ∼5,000 years ago (*9*), it is important to note that humans’ unique ability for language is substantially older, emerging at least 135,000 years ago (*47, 48*). Moreover, humans started producing abstract engravings almost 100,000 years ago (*10*) and hence in a timeframe during which the human brain underwent substantial evolutionary changes (*11*). While it is debated whether the earliest engravings were used for communication (*49, 50*), or merely for decoration (*51*), figurative depictions appeared at least 50,000 years ago, suggesting a long history of cultural engagement with symbols (*52, 53*). As such, while formal writing systems gradually abstracted away from their iconographic roots (*12, 54*), repeated engagement may have shaped neural systems linking vision and language through bio-cultural feedback loops (*13*).

If humans indeed have a predisposition for vision-language mapping, an ability to extract language content from visual input should be detectable early on in development. Indeed there is evidence that children become “symbol-minded”, i.e. understand that one thing represents something other than itself, remarkably early in development (*55, 56*). Even before acquiring spoken language, infants can recognize that a two-dimensional picture is the visual representation of a real-world object (*14*) and they can even learn to use visual “baby signs” for communication (*57*). Infants are also already able to match visual and auditory speech information across modalities (*58*). Together with our data, these findings may suggest that reading text represents a highly abstract extension of an innate vision-language mapping capacity that unfolds developmentally, progressing from early gestures (*59*) to word-object mappings (*60*), to understanding abstract visual symbols as representations of the world (*61, 62*). Future studies could further explore the role of mOTS-words in vision-language mapping across development, such as the region’s engagement in processing baby signs during infancy.

Our results may have important implications for developmental dyslexia, as altered structural (*63*) and functional (*64, 65*) properties of OTS-words represent a key characteristic of this reading disorder. Our data suggest that these alterations may result from a reduced engagement of mOTS-words in symbolic language processing that precedes literacy training. In accordance with this proposal, dyslexia has been associated with changes in functional and structural connectivity between OTS-words and the language network (*66, 67*), which are shaped in part by genetic factors (*22, 68, 69*). Critically, our paradigm may be suitable to detect an insufficient engagement of OTS-words before the onset of reading instruction, raising the possibility that symbolic language processing prior to schooling could serve as a biomarker to facilitate early diagnosis and targeted interventions during a developmental window when reading-related neural circuits remain most plastic (*70*). These interventions could focus on training symbol-meaning mapping to increase OTS-words symbolic language engagement prior to the onset of literacy training.

Together, these findings suggest that learning to read builds on a neural scaffold that links vision and language with broad implications for cultural anthropology, developmental science, education, and clinical neuroscience.

## Funding

European Union (ERC, WRAPPED, 101161197). Views and opinions expressed are however those of the author(s) only and do not necessarily reflect those of the European Union or the European Research Council. Neither the European Union nor the granting authority can be held responsible for them.

Deutsche Forschungsgemeinschaft (German Research Foundation, DFG) under Germany’s Excellence Strategy (EXC 3066/1 “The Adaptive Mind”, Project No. 533717223)

Deutsche Forschungsgemeinschaft (DFG, German Research Foundation), project number 222641018 – SFB/TRR 135 TP C10

LOEWE Professorship awarded by the State of Hesse (LOEWE/4b//519/05/01.002(0016)/120)

Lise Meitner Excellence Program of the Max Planck Society

## Author contributions

Conceptualization: MG, AS, AD Methodology: MG, AD, MP, AU, AS

Investigation: AD, MP, AU, AS, MK, MM, LF, VAAG, GC,

Visualization: AD, AU, AS

Funding acquisition: MG

Project administration: MG, AD

Supervision: MG, GS, BDH, YLS

Writing – original draft: MG, AD

Writing – review & editing: MG, AD, BDH

Authors declare that they have no competing interests.

## Data, code, and materials availability

Data was processed using open-source software including FreeSurfer (https://surfer.nmr.mgh.harvard.edu/) and mrVista (https://github.com/vistalab/vistasoft). Custom analysis code and data necessary to reproduce all figures in the main manuscript are available at https://github.com/EduNeuroLab/read_emojis_kids.

