## Supplementary Material for "Preliterate symbolic language processing sets the neural stage for learning to read"

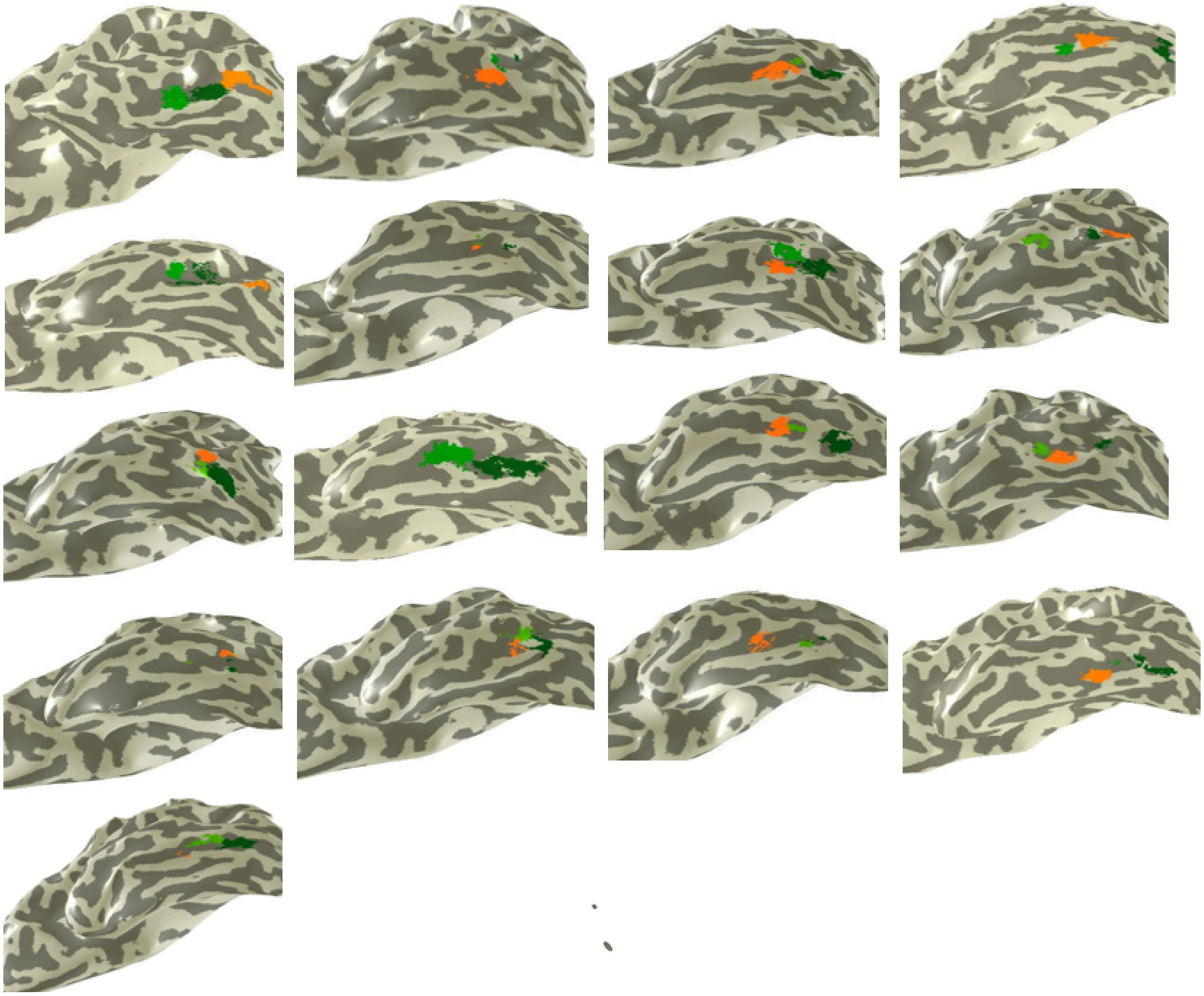

**Fig. S1. Functional regions of interest (ROIs) were defined in each individual participant**. Three functional ROIs - mOTS-words (light green), pOTS-words (dark green), and OTS-limbs (orange) - are shown for each individual child on their left inflated cortical surface. ROIs were defined in the occipito-temporal cortex (OTS) from experiment 1 data collected after the children completed first grade by contrasting words with the other categories (OTS-words subregions; T≥3, voxel level, uncorrected) or by contrasting limbs with the other categories (OTS-limbs; T≥3, voxel level, uncorrected).

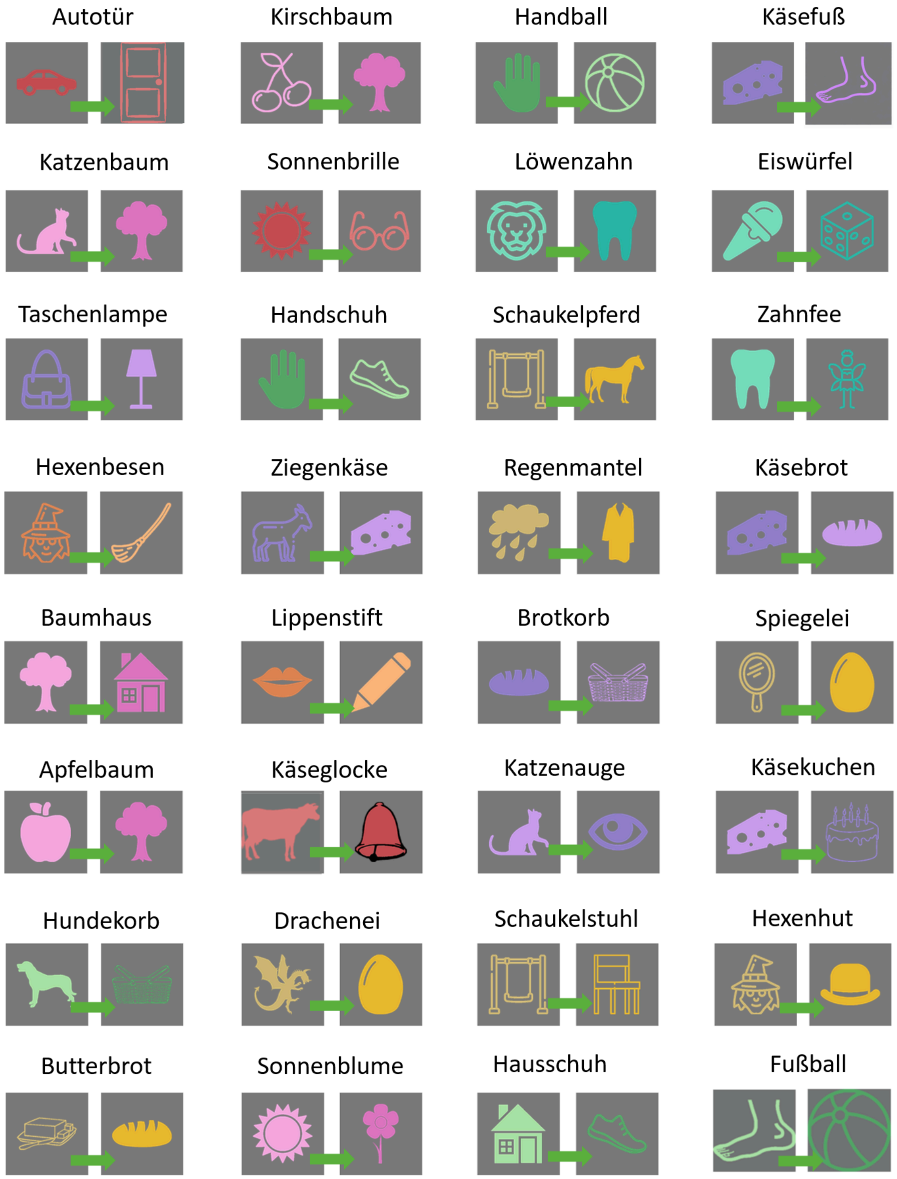


**Fig. S2 Overview of symbolic compound nouns used in experiment 2.** Stimuli were German compound nouns, presented as symbols, and were shown in the same or slightly different color hues (matching or mismatching). Stimuli either formed a meaningful compound (Baum+Haus ⟶ "tree house") or were shown in reversed order (Haus+Baum ⟶ "house tree") and thereby formed a meaningless compound. Participants performed either a symbolic language task (meaningful or meaningless) or a color task (same hue or different hue) on these stimuli. List of compound meaning from left to right: car door, cherry tree, hand ball, cheesy foot, cat tree, sun glasses, dandelion, ice cube, torch, glove, rocking horse, tooth fairy, witches' broom, goat cheese, rain coat, cheese bread, tree house, lipstick, bread basket, sunny-side-up, apple tree, cheese bell, cat eye, cheese cake, dog basket, dragon egg, rocking chair, witches hat, buttered bread, sunflower, house shoe, football

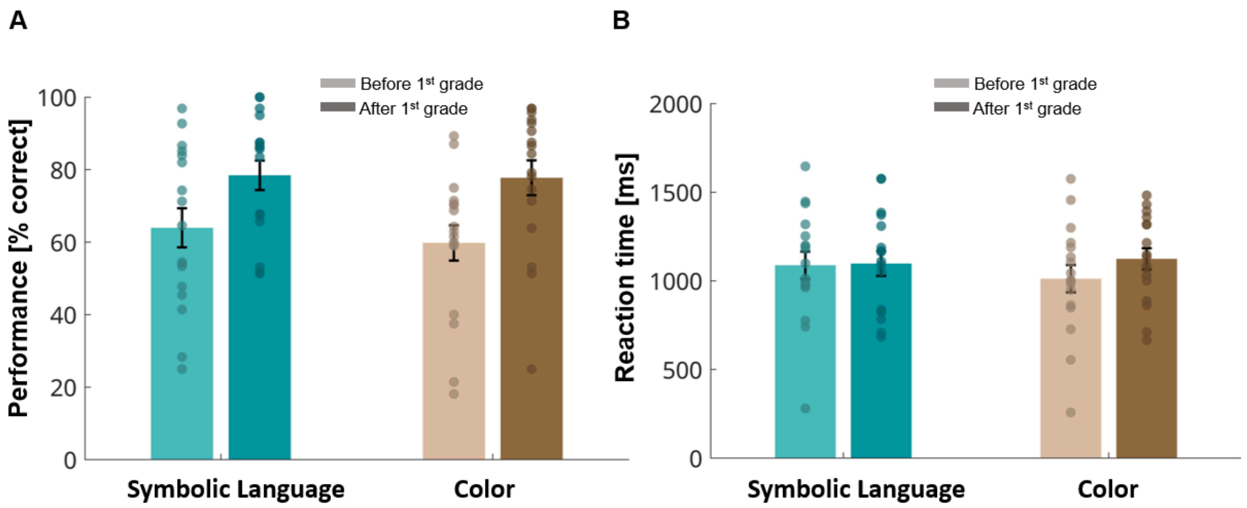


**Fig S3. Behavioral responses in fMRI Experiment 2.** (A) Performance (percentage of trials with an accurate response) for symbolic language and color task before (light bars) and after (dark bars) the first year of schooling (n = 17). Accuracy exceeded chance level (50%) for both tasks and timepoints (symbolic language: before first grade: 64±5%, $t(16)=2.58$, $p=0.02$, after first grade: 78±4%, $t(16)=6.98$, $p<0.0001$; color task: before first grade: 60±5%, $t(16)=2.27$, $p=0.04$, after first grade: 78±5%, $t(16)=5.79$, $p<0.0001$). Performance increased for both tasks after first grade (main effect of time, $F(1,16)=7.96$, $p=0.01$), and there was no main effect of task ($F(1,16)=0.40$, $p=0.54$) and no task × time interaction ($F(1,16)=0.73$, $p=0.41$). (B) Reaction times for linguistic and color conditions before (light bars) and after (dark bars) the first year of schooling. Repeated measures ANOVAs revealed no change in reaction times over the first year of schooling (no main effect of time: $F(1,16)=0.58$, $p=0.45$), and no significant difference between tasks at either timepoint (no main effect of task $F(1,16)=0.39$, $p=0.54$ no task x time interaction $F(1,16)=2.60$, $p=0.12$). Bars represent group means ± SEM; dots indicate individual participant data.

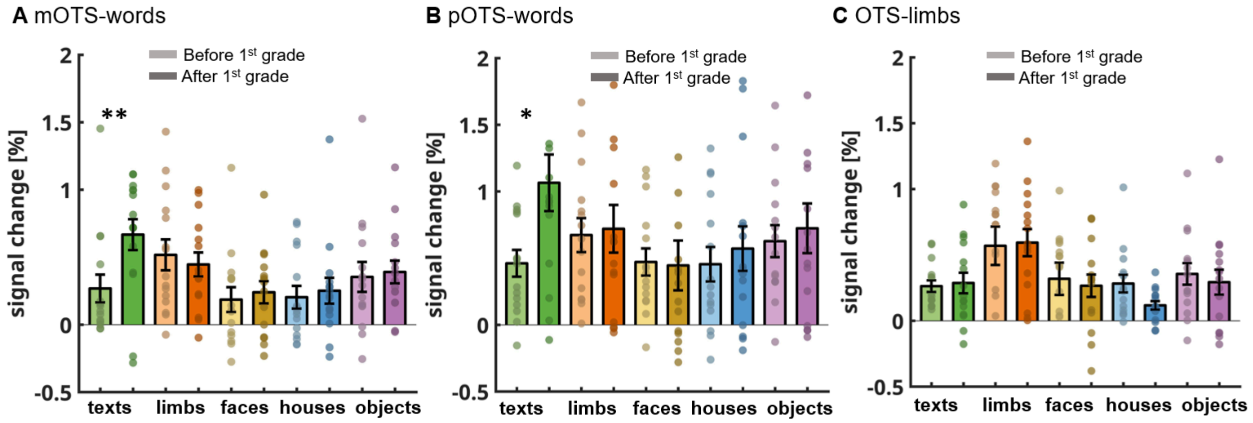


**Fig. S4. OTS-words subregions show increasing responses to texts during the first year of schooling.** (A–C) Mean percentage signal change (± SEM) for five visual categories (texts, limbs, faces, houses, and objects) are shown before (lighter shades) and after (darker shades) first grade in three ROIs: mOTS-words (A), pOTS-words (B), and OTS-limbs (C). (A) mOTS-words showed a significant main effect of category ($F(4,11)=7.63$, $p<0.0001$) and a significant time × category interaction ($F(4,11)=8.20$, $p<0.0001$), with no significant main effect of time ($F(1,14)=1.36$, $p=0.26$). Responses to text increased significantly during the first year of schooling ($p=0.007$), with no other category changing significantly (all ps>0.06). Before school, limbs elicited the highest responses (higher than texts: $p=0.04$; faces: $p=0.004$); after first grade, texts elicited the highest responses (higher than limbs: $p=0.001$, faces: $p<0.0001$, houses: $p=0.03$, objects: $p=0.01$). (B) pOTS-words showed a significant main effect of category ($F(4,11)=10.98$, $p<0.0001$) and a significant time × category interaction ($F(4,11)=16.05$, $p<0.0001$), with no main effect of time ($F(1,14)=0.76$, $p=0.39$). Responses to text increased significantly during first grade ($p=0.01$), with no other category changing significantly (all $p>0.58$). Before school, responses were not higher for any given category over any other (all ps>0.05); after first grade, texts elicited higher responses than all other categories (limbs: $p=0.003$, faces: $p<0.0001$, houses: $p<0.0001$, objects: $p=0.007$). (C) OTS-limbs showed a main effect of category ($F(4,11)=8.59$, $p<0.0001$, limbs higher than texts: $p=0.005$, faces: $p=0.003$, objects: $p=0.04$), no main effect of time ($F(1,14)=0.87$, $p=0.37$) and no time × category interaction ($F(4,11)=0.93$, $p=0.45$). Asterisks are based on post-hoc analyses and indicate a significant change in response to a given category during the first year of schooling (** $p<0.01$; * $p<0.05$).

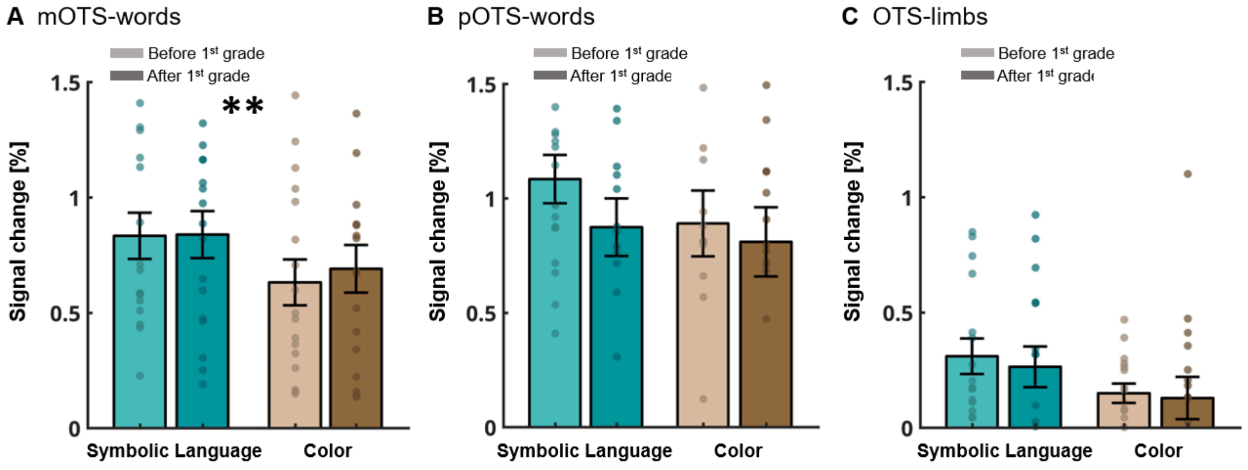


**Fig. S5. mOTS-words is engaged in symbolic language processing both before and after first grade.** (A–C) Mean percentage signal change (± SEM) for the symbolic language and color tasks in experiment 2 are shown before (lighter shades) and after (darker shades) first grade in three ROIs: mOTS-words (A), pOTS-words (B), and OTS-limbs (C). (A) mOTS-words shows a main effect of task ($F(1,16)=11.34$, $p=0.004$), with no main effect of time ($F(1,16)=0.10$, $p=0.76$) and no significant time × task interaction ($F(1,16)=1.49$, $p=0.24$). Responses were higher in the symbolic language task than the color task both before ($p=0.008$) and after ($p=0.004$) first grade. (B-C) pOTS-words (B) and OTS-limbs (C) showed no main effects of task, time, or task × time interactions (all ps>0.10). Circles show individual subject data, asterisks indicate main effect of task (** $p < 0.01$).

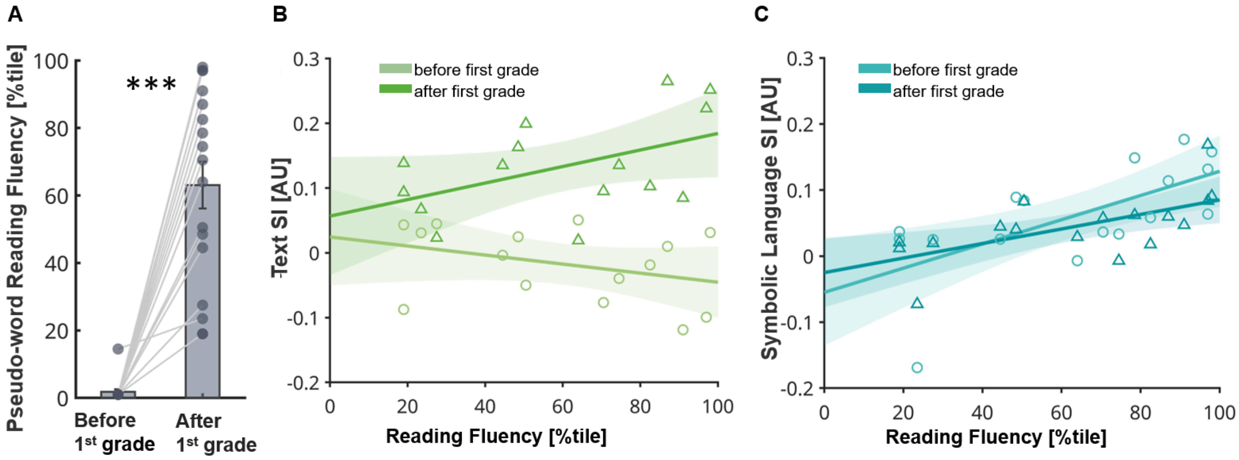


**Fig. S6. Preliterate symbolic language processing in mOTS-words predicts future pseudo-word reading fluency.** (A) Pseudo-word reading fluency (SLRT-II, percentile) increased during the first year of schooling (paired t-test: $t(16)=8.44$, $p<0.0001$). Bars show mean percentile relative to a normative sample (± SEM), each circle is an individual, lines connect the two measurements of the same individual. (B) Individual children's (N=15) pseudo-word reading fluency after first grade did not correlate with mOTS-words text selectivity before first grade (circles, light green; $r^2=0.12$, $p=0.20$) and it only marginally correlated with mOTS-words text selectivity after first grade (triangles, dark green; $r^2=0.23$, $p=0.07$). (C) Individual children's (N=17) pseudo-word reading fluency after first grade correlates with mOTS-words symbolic language selectivity after first grade (triangles, dark cyan; $r^2=0.39$, $p=0.008$) and is predicted by mOTS-words symbolic language selectivity before first grade (circles, light cyan; $r^2=0.43$, $p=0.004$). In (B,C) each triangle or circle is an individual, lines show least-squares regression fits, shaded regions show 95% confidence intervals.
